# Determination of effective meropenem and gentamicin doses in a silkworm infection model using a clinical *Klebsiella aerogenes* isolate

**DOI:** 10.64898/2026.08.25.746979

**Authors:** Shinobu Hirayama, Yasuhiko Matsumoto, Sanae Kurakado, Mariko Otani, Takahiro Matsumoto, Hinako Murakami, Kazuhiro Tateda, Takashi Sugita

## Abstract

*Klebsiella aerogenes*, a member of the Enterobacteriaceae, is a causative agent of healthcare-associated infections, and outbreaks caused by drug-resistant *K. aerogenes* have been reported worldwide. The range of antimicrobial agents available for treating infections caused by carbapenem-resistant *K. aerogenes* is limited. While *in vivo* animal experiments using clinical *K. aerogenes* isolates to evaluate antimicrobial therapy could facilitate selection of the most effective treatment, conducting infection experiments involving large numbers of mammals such as mice is challenging due to ethical concerns related to animal welfare. Silkworms are invertebrates increasingly used as experimental models for infectious disease research to evaluate antimicrobial efficacy. In this study, we aimed to establish a silkworm infection model using a clinical *K. aerogenes* isolate to evaluate its utility for determining effective antimicrobial doses. *K. aerogenes* strains were isolated from a patient at a Japanese hospital, and a silkworm infection model was established using the clinical isolate. The non-metallo-beta-lactamase-producing strain *K. aerogenes* TUM25562, isolated from a patient with a complicated urinary tract infection, was susceptible to meropenem (MEPM) and gentamicin (GM) *in vitro*. During treatment, additional isolates with increased resistance to MEPM and subsequently to both MEPM and GM emerged. *K. aerogenes* TUM25562 caused dose-dependent mortality in silkworms. Treatment with clinically equivalent weight-based doses of MEPM or GM did not cure the infected silkworms. The median effective (ED_50_) doses of MEPM and GM were therefore investigated using the silkworm infection model. Administration of higher doses corresponding to four times the ED_50_ significantly prolonged the survival of infected silkworms. These results suggest that a silkworm infection model using clinical *K. aerogenes* isolates may provide a practical approach for evaluating antimicrobial efficacy and determining effective antimicrobial doses.

## Introduction

*Klebsiella aerogenes*, a pathogenic bacterium belonging to the Enterobacteriaceae, causes urinary tract infections, pneumonia, and sepsis [1,2]. In recent years, clinical cases of nosocomial infection caused by *K. aerogenes* have been reported worldwide [1,3–5]. Several *K. aerogenes* isolates from patients exhibit antimicrobial resistance mediated by mechanisms such as carbapenemase, AmpC, or extended-spectrum beta-lactamase production, as well as efflux pump or porin mutations [6–9].

The IDSA 2024 guidance on the treatment of antimicrobial-resistant Gram-negative infections recommends carbapenems such as meropenem (MEPM) for the treatment of infections caused by carbapenemase-nonproducing and AmpC-producing Enterobacterales because carbapenems are stable to AmpC-mediated hydrolysis [10]. MEPM is also recommended for the treatment of *K. aerogenes* strains resistant to β-lactam antibacterial drugs [11]. Furthermore, when drug susceptibility has been confirmed and there is a risk of resistance or toxicity associated with sulfamethoxazole-trimethoprim or fluoroquinolones, cefepime (CFPM) or aminoglycosides are recommended as alternative therapeutic agents [10]. Although the meta-analysis cited in the guidance found no significant difference in clinical outcomes between patients treated with CFPM and those treated with carbapenems, carbapenems are generally selected for patients with severe infections [10,11]. Similarly, a cohort study comparing CFPM and MEPM for bacteremia caused by Enterobacteriaceae with a moderate risk of AmpC production, including *K. aerogenes*, reported comparable 90-day mortality rates (18.9% vs 17.1%) [12–14]. Moreover, *K. aerogenes* bacteremia has been associated with poorer clinical outcomes than *Enterobacter cloacae* complex bacteremia, including higher rates of in-hospital mortality, recurrent bacteremia, and complications [15]. Consequently, treating severe *K. aerogenes* infections remains challenging, and selecting the most appropriate antimicrobial agent can be difficult. Although *in vitro* antimicrobial susceptibility testing provides valuable guidance, it does not always identify the optimal treatment regimen. An *in vivo* infection model capable of evaluating antimicrobial efficacy and safety using clinical isolates could complement susceptibility testing by providing additional information on therapeutic efficacy and safety. Conducting infection experiments involving large numbers of mammals such as mice, however, is challenging due to ethical concerns related to animal welfare.

Silkworms, invertebrate animals, are increasingly used as experimental models for infectious disease research [16–18]. Because silkworms raise fewer animal welfare concerns than mammals such as mice, they offer advantages for conducting infection experiments involving large numbers of animals [17, 19]. Silkworm infection models have been established to evaluate the pathogenicity of human pathogens, including *Staphylococcus aureus* [17,18]. In addition, virulence identified using silkworm infection models correlates with pathogenicity in mice [20, 21]. These findings support the use of silkworm infection models for investigating bacterial pathogens that infect mammals.

Silkworm infection models are also useful for quantitatively evaluating the therapeutic efficacy of antimicrobial agents [22,23]. Antimicrobial efficacy can be quantitatively assessed by determining the median effective dose (ED_50_), defined as the dose required for half of the infected silkworms to survive [24]. Similarly, compound toxicity can be evaluated by determining the half maximal lethal dose (LD_50_), and the LD_50_ values of toxic compounds in silkworms correlate with those reported in mammals [25]. Together, these findings demonstrate that silkworm infection models provide a quantitative platform for evaluating both therapeutic efficacy and toxicity [26,27]. Accordingly, *in vivo* chemical screening using silkworm infection models has identified compounds with therapeutic efficacy in mouse infection models of *S. aureus* and *Aspergillus fumigatus* [28–30]. These findings suggest that silkworm infection models can be used to evaluate compounds with pharmacologic activity in mammals. Recently, a silkworm infection model was established for *K. aerogenes* to evaluate bacterial pathogenicity and the pharmacologic effects of various compounds [31]. The potential of such a model to simulate antimicrobial treatment using clinical *K. aerogenes* isolates and support antimicrobial selection, however, has not been investigated.

In this study, we established a silkworm infection model using a *K. aerogenes* strain isolated from a patient with a difficult-to-treat infection. We then evaluated the therapeutic efficacy of antimicrobial agents used in clinical practice and investigated whether the model could be used to determine effective antimicrobial doses. Our findings demonstrate the potential of a silkworm infection model using clinical isolates as a practical *in vivo* platform to complement antimicrobial susceptibility testing and support antimicrobial selection.

## Materials and Methods

### Ethics approval, clinical specimen, and bacterial isolation

This study was approved by the Ethics Committee of the Toho University School of Medicine (Approval Nos. A24108, A24045, and A23052) and authorized by Meiji Pharmaceutical University (Approval No. R5-006).

### Bacterial isolation from a patient

Bacterial isolation was performed as described previously [32]. A clinical urine specimen (1 µL) was inoculated onto Accurate™ Fractionated Sheep Blood Agar/McConkey Medium (Shimadzu Diagnostics) and incubated at 35°C under CO_2_-enriched conditions for 18 h. Colonies presumptively identified as members of the order Enterobacterales were collected, and bacterial suspensions were prepared using the Prompt method. The bacterial suspensions were dispensed onto NC-EN2T panels using the AutoLynoc BRID® system (Beckman Coulter), and bacterial identification and antimicrobial susceptibility testing were performed using the MicroScan WalkAway system (Beckman Coulter). *Escherichia coli* ATCC 25922 was used for quality control.

Carbapenemase production was investigated in isolates with an MEPM MIC of ≥2 µg/mL, a tazobactam/piperacillin MIC of ≥16/4 μg/mL, or a latamoxef MIC of >8 μg/mL. The modified carbapenem inactivation method (mCIM) was performed first. Isolates positive by the mCIM were subsequently examined using the sodium mercaptoacetic acid disk method. Isolates positive by both the mCIM method and the sodium mercaptoacetic acid disk method were classified as metallo-β-lactamase-producing strains. Isolates negative by the mCIM were classified as non-carbapenemase-producing strains.

### Reagents

MEPM trihydrate, GM sulfate, NaCl, and agar were purchased from Fujifilm Wako Pure Chemical Industries (Osaka, Japan). Amikacin (AMK) sulfate was obtained from Tokyo Chemical Industry Co., Ltd. (Tokyo, Japan). Tryptone and yeast extract were purchased from Becton, Dickinson and Company (Franklin Lakes, NJ, USA). Cefepime was purchased from MedChemExpress (Monmouth Junction, NJ, USA).

### *K. aerogenes* culture conditions

*K. aerogenes* strains (TUM25562; isolation date: 20/10/2021, TUM25564; isolation date: 18/12/2021, and TUM25556; isolation date: 4/2/2022) were cultured on Luria Bertani (LB) agar (1% tryptone, 0.5% yeast extract, 1% NaCl, and 1.5% agar) and incubated at 37°C for 24 h.

### Antimicrobial susceptibility test

Antimicrobial susceptibility testing was performed using the microdilution method with Eiken DP41 dry plates (Eiken Chemical Co., Ltd., Tokyo, Japan) and Eiken Mueller-Hinton broth (Eiken Chemical Co., Ltd., Tokyo, Japan). The results were interpreted according to the CLSI M100 breakpoints. Each *K. aerogenes* strain was suspended in sterile saline, and the suspension was adjusted to an optical density at 600 nm (OD_600_) of 0.1. An aliquot of the adjusted *K. aerogenes* cell suspension (25 µL) was added to 12 mL of Eiken Mueller-Hinton broth. The resulting bacterial suspension was inoculated into each well of a dry plate in 100-µL aliquots. The final inoculum was approximately 5 × 10^5^ CFU/well. The plates were incubated at 37°C for 16–20 h. Bacterial growth was assessed visually. Growth was considered positive when turbidity or a precipitate with a diameter of >1 mm was observed or when two or more precipitate clumps were observed, even if each clump was <1 mm in diameter.

### Silkworm infection experiments

Silkworm infection experiments were performed as described previously [33,34]. Silkworm eggs (KINSYU × SHOWA) were purchased from Ehime-Sanshu Co., Ltd. (Ehime, Japan). Fifth-instar larvae were fed overnight with an artificial diet before use (Silkmate 2S; Ehime-Sanshu Co., Ltd.). *K. aerogenes* strains were cultured on LB agar at 37°C for 24 h. *K. aerogenes* cells were collected from agar plates, suspended in saline, and adjusted to an OD_600_ of 4. Each *K. aerogenes* cell suspension (50 µL) was injected into the silkworm hemolymph from the dorsal surface using a 1-mL tuberculin syringe (Terumo Medical Corporation, Tokyo, Japan). The infected silkworms were maintained at 37°C and monitored for 3 days. Survival curves were generated using the Kaplan–Meier method and compared using the log-rank (Mantel–Cox) test in GraphPad Prism version 11.0.2 (GraphPad Software, LLC, San Diego, CA, USA).

### LD_50_ determination in the silkworm infection model

The LD_50_ of *K. aerogenes* was determined by modifying a previously reported method [35,36]. The bacterial suspension was serially diluted twofold with saline. Each silkworm was injected with 2.6 × 10^7^ to 4.1 × 10^8^ bacterial cells in a 50-µL volume) and maintained at 37°C. Silkworm survival (n = 4 per group) was assessed 24 h after injection. The LD_50_ was determined by simple logistic regression using combined data from two independent experiments in GraphPad Prism software version 11.0.2.

### Viable counts of *K. aerogenes* in the hemolymph of infected silkworms

Viable bacterial counts in hemolymph of the infected silkworms were determined by modifying a previously reported method [37]. Silkworms were injected with a *K. aerogenes* TUM 25562 cell suspension (7 x 10^6^ cells in 50 µl) and incubated at 37°C. Hemolymph was harvested from the silkworm larvae through a cut in the first proleg at 1, 3, or 6 h after infection. The hemolymph was diluted in saline and spread onto an LB agar plate. After incubation at 37°C for 24 h, the colonies were counted, and viable bacterial counts were determined.

### Evaluation of antimicrobial efficacy in the silkworm infection model

The therapeutic efficacy of antimicrobial agents was determined using the silkworm infection model by modifying a previously reported method [38]. Silkworms were injected with a *K. aerogenes* TUM 25562 cell suspension (2–4 x 10^8^ cells/50 µl). For the experiment shown in Figure 5, MEPM solution (80 µg/50 µL) or saline (50 µL) was administered at 2, 6, and 10 h after infection. For the experiment shown in Figure 6, GM solution (15 µg/50 µL), AMK solution (30 µg/50 µL), or saline (50 µL) was administered at 2 h after infection. For the experiment shown in Figure 7, CFPM solution (100 µg/50 µL) or saline (50 µL) was administered at 2, 4, and 6 h after infection. For the experiments shown in Figures 9 and 10, MEPM solution (500 µg/50 µL), GM solution (800 µg/50 µL), or saline (50 µL) was administered at 2 h after infection. The infected silkworms were incubated at 37°C and monitored for survival for 3 days. Survival curves were generated using the Kaplan–Meier method and compared using the log-rank (Mantel–Cox) test in GraphPad Prism version 11.0.2.

### ED_50_ determination in the silkworm infection model

The ED_50_ of each antimicrobial agent was determined using the silkworm infection model by modifying a previously reported method [39]. Silkworms were injected with *K. aerogenes* cells (2–4 × 10^8^ cells/50 µL). At 2 h after infection, each silkworm was injected with various concentrations of the antimicrobial agents (50 μL) dissolved in saline. The antimicrobial agents were tested at doses generated by twofold serial dilution. Five or six silkworms were used for each dose, and survival was assessed at 24 h after infection. ED_50_ values were estimated by simple logistic regression using combined data from three independent experiments in GraphPad Prism software version 11.0.2.

### Statistical analysis

All experiments were performed at least three times, and representative results are presented. Statistical significance between the saline group and the drug-treated group was determined using the log-rank (Mantel–Cox) test. *P* < 0.05 was considered statistically significant. Statistical analyses were performed using GraphPad Prism version 11.0.2.

## Results

### Isolation of *K. aerogenes* strains from a patient

The patient was a 1-year-old child born via cesarean section at 37 weeks and 2 days of gestation (height: 59.7 cm; weight: 4000 g). The patient had trisomy 18, vermis hypoplasia, bilateral great vessels arising from the right ventricle, subaortic ventricular septal defect, and esophageal atresia. In accordance with the patient’s clinical condition and growth, pulmonary artery banding, radical esophageal atresia repair, and laryngotracheal separation were performed. Since birth, the patient has experienced recurrent episodes of aspiration pneumonia, urinary tract infections, and catheter-related infection requiring antimicrobial therapy. Treatment with cefotaxime (CTX) was initiated (Fig. 1). In October 2021, elevated inflammatory markers led to suspicion of aspiration pneumonia, a urinary tract infection, and a catheter-related infection. After treatment with CTX, *K. aerogenes* TUM25562 was isolated from a urine specimen. Because the inflammatory markers showed little improvement and a thoracic drain infection was suspected, CTX was switched to MEPM. Approximately 2 months after the initiation of CTX and subsequent MEPM therapy, *K. aerogenes* TUM25564 was isolated from a urine specimen. Treatment with MEPM followed by GM was continued for an additional 1.5 months. In February 2022, a third isolate, *K. aerogenes* TUM25556, was recovered from a urine specimen (Fig. 1). Antimicrobial susceptibility testing using the MicroScan WalkAway system was performed on all three isolates (TUM25562, TUM25564, and TUM25556) (Table S1). None of the three *K. aerogenes* isolates produced carbapenemase (Table S2).

**Fig. 1.**
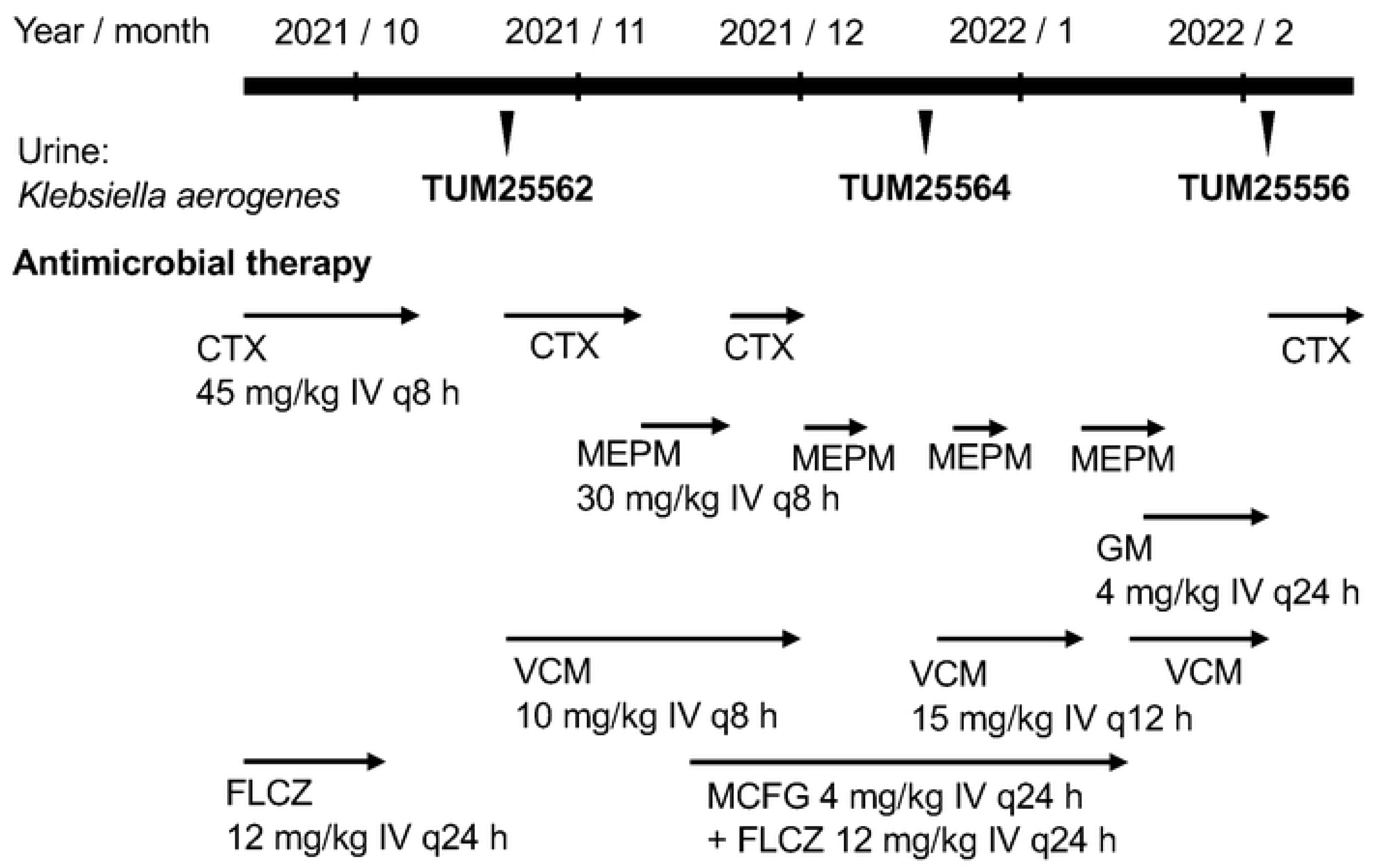
Timeline of antimicrobial therapy and *K. aerogenes* isolation in a patient. A 1-year-old patient (height: 59.7 cm, weight: 4000 g) born via cesarean section at 37 weeks and 2 days of gestation. In October 2021, elevated inflammatory markers led to suspicion of aspiration pneumonia, a urinary tract infection, and a catheter-related infection. After treatment with cefotaxime (CTX), *K. aerogenes* (TUM 25562) was isolated from a urine specimen. Approximately 2 months after administering CTX or MEPM, *K. aerogenes* (TUM 25564) was isolated from a urine specimen. MEPM or GM was administered for an additional 1.5 months, and in February 2022, *K. aerogenes* (TUM25556) was isolated.

### Drug sensitivity of *K. aerogenes* strains

The antimicrobial susceptibility of *K. aerogenes* strains TUM25562, TUM25564, and TUM25556 was evaluated by determining MIC values of MEPM, GM, AMK, and CFPM based on the CLSI criteria (Table 1). TUM25562 was susceptible to all four antimicrobial agents, with MIC values of 0.5, ≤2, ≤8, and 4 for MEPM, GM, AMK, and CFPM, respectively (Table 1). TUM25564 showed reduced susceptibility to MEPM and CFPM, with MIC values of 8, ≤2, 16, and 8 µg/mL, respectively (Table 1). TUM25556 exhibited resistance to both MEPM and GM, with MIC values of >8, while remaining susceptible to AMK (MIC ≤8 µg/mL); the MIC of CFPM was >16 µg/mL (Table 1).

**Table 1.** MIC values of antimicrobial agents against clinical *K. aerogenes* isolates.

| Antibacterial drug | TUM25562 | TUM25564 | TUM25556 |
| --- | --- | --- | --- |
| Piperacillin | > 64 | > 64 | > 64 |
| Ampicillin | > 16 | > 16 | > 16 |
| Fosfomycin | ≤ 32 | 128 | 128 |
| Tazobactam/Piperacillin | 4/64 | 4/64 | > 4/64 |
| Sulbactam/Ampicillin | > 8/16 | > 8/16 | > 8/16 |
| Minocycline | 4 | > 8 | 8 |
| Cefazolin | > 16 | > 16 | > 16 |
| Ceftriaxone | 32 | > 32 | > 32 |
| Ceftazidime | > 16 | > 16 | > 16 |
| Cefmetazole | > 32 | > 32 | > 32 |
| Cefpodoxime | > 4 | > 4 | > 4 |
| Gentamicin | ≤ 2 | ≤ 2 | > 8 |
| Amikacin | ≤ 8 | 16 | ≤ 8 |
| Sulfamethoxazole-Trimethoprim | > 38/2 | > 38/2 | > 38/2 |
| Cefotiam | > 4 | > 4 | > 4 |
| Cefepime | 4 | 8 | > 16 |
| Flomoxef | > 16 | > 16 | > 16 |
| Imipenem | 2 | 8 | > 8 |
| Levofloxacin | 0.25 | 0.5 | 0.5 |
| Meropenem | 0.5 | 8 | > 8 |
| Aztreonam | 16 | > 16 | > 16 |

### Establishment of a silkworm infection model using a *K. aerogenes* clinical isolate TUM25562

A silkworm infection model using a multi-drug-resistant *S. aureus* strain was previously established to evaluate the *in vivo* efficacy of antimicrobial agents [40,41]. In the present study, we established a silkworm infection model using the clinical *K. aerogenes* isolate TUM25562. Injection of *K. aerogenes* TUM25562 (1 × 10⁸ cells/silkworm) resulted in the death of four of five silkworms within 48 h (Fig. 2A). To evaluate the dose-dependent lethality of the isolate, silkworms were injected with serially diluted bacterial suspensions (Fig. 2B). Mortality increased in a dose-dependent manner, and the LD_50_ value of *K. aerogenes* TUM25562 was determined from the survival data (Fig. 2B, Table 2). Using the same experimental system, the LD_50_ values of *K. aerogenes* TUM25564 and TUM25556 were also determined (Table 2, Fig. S1). To evaluate whether *K. aerogenes* TUM25562 proliferated *in vivo*, viable bacterial counts in the hemolymph of the infected silkworms were measured. The number of viable *K. aerogenes* TUM25562 cells increased during the first 3 h after infection (Fig. 3). Next, we investigated whether bacterial viability was required for silkworm death. Silkworms injected with autoclave-treated *K. aerogenes* TUM25562 cells survived for at least 2 days after injection (Fig. 4). Together, these results demonstrate that *K. aerogenes* TUM25562 establishes a lethal, proliferative infection in silkworms and validate this model for subsequent evaluation of antimicrobial efficacy.

**Fig. 2.**
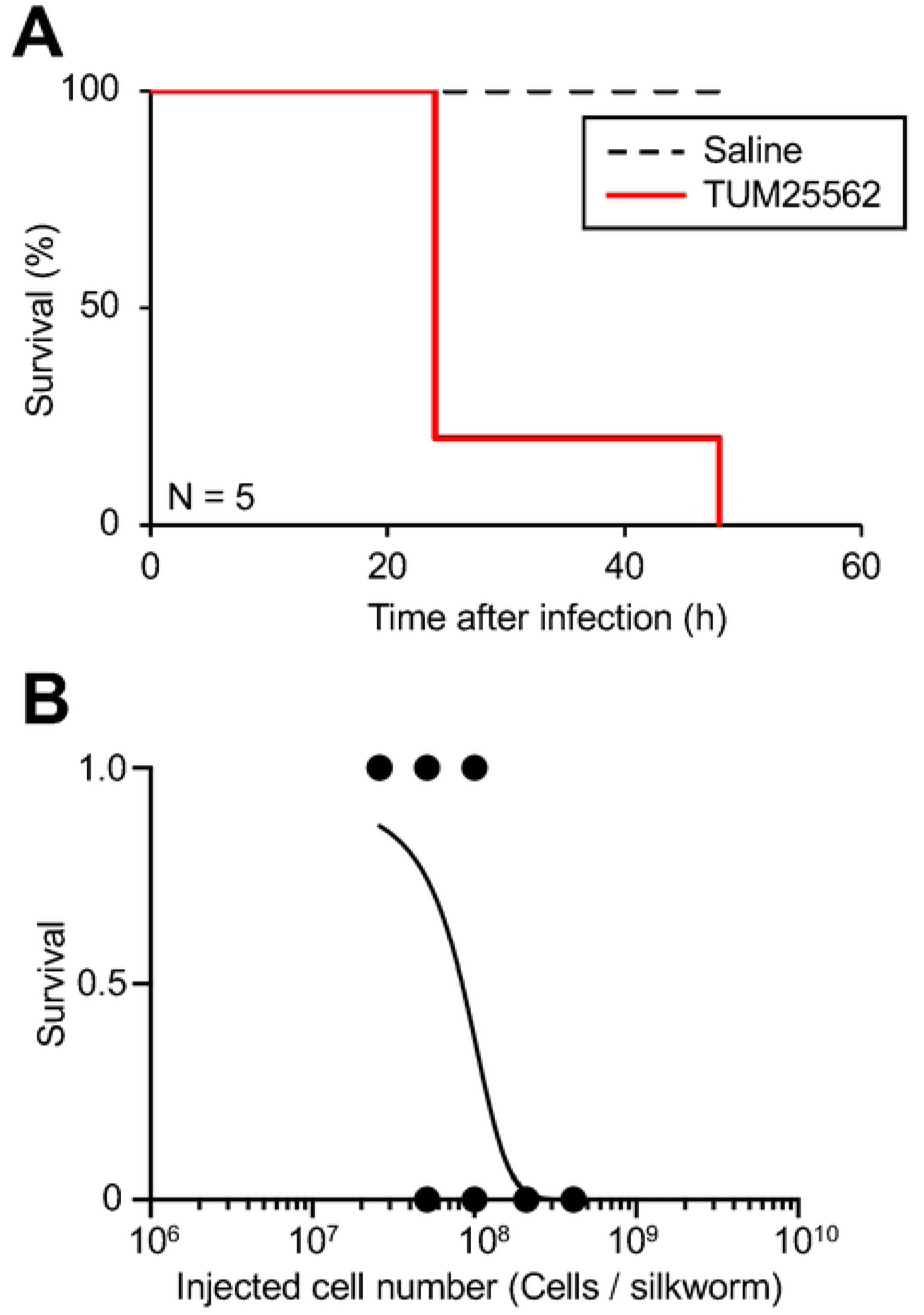
Silkworm killing ability of *K. aerogenes* TUM25562. (**A**) Time course of survival of silkworms injected with *K. aerogenes* TUM25562. The *K. aerogenes* TUM25562 suspension (1 × 10⁸ cells/silkworm) or saline was injected into the silkworm hemolymph. After injection, the silkworms were incubated at 37°C for 2 days. Survival curves were generated using the Kaplan–Meier method. N = 5/group. (**B**) Dose-dependency of *K. aerogenes* TUM25562 on silkworm survival. The survival rates of silkworms at 37°C were determined 24 h after *K. aerogenes* TUM25562 injection. Surviving and dead silkworms were scored as 1 and 0, respectively. A total of 40 silkworms were used per group. Curves were drawn by combining the results of two independent experiments using a simple logistic regression model.

**Fig. 3.**
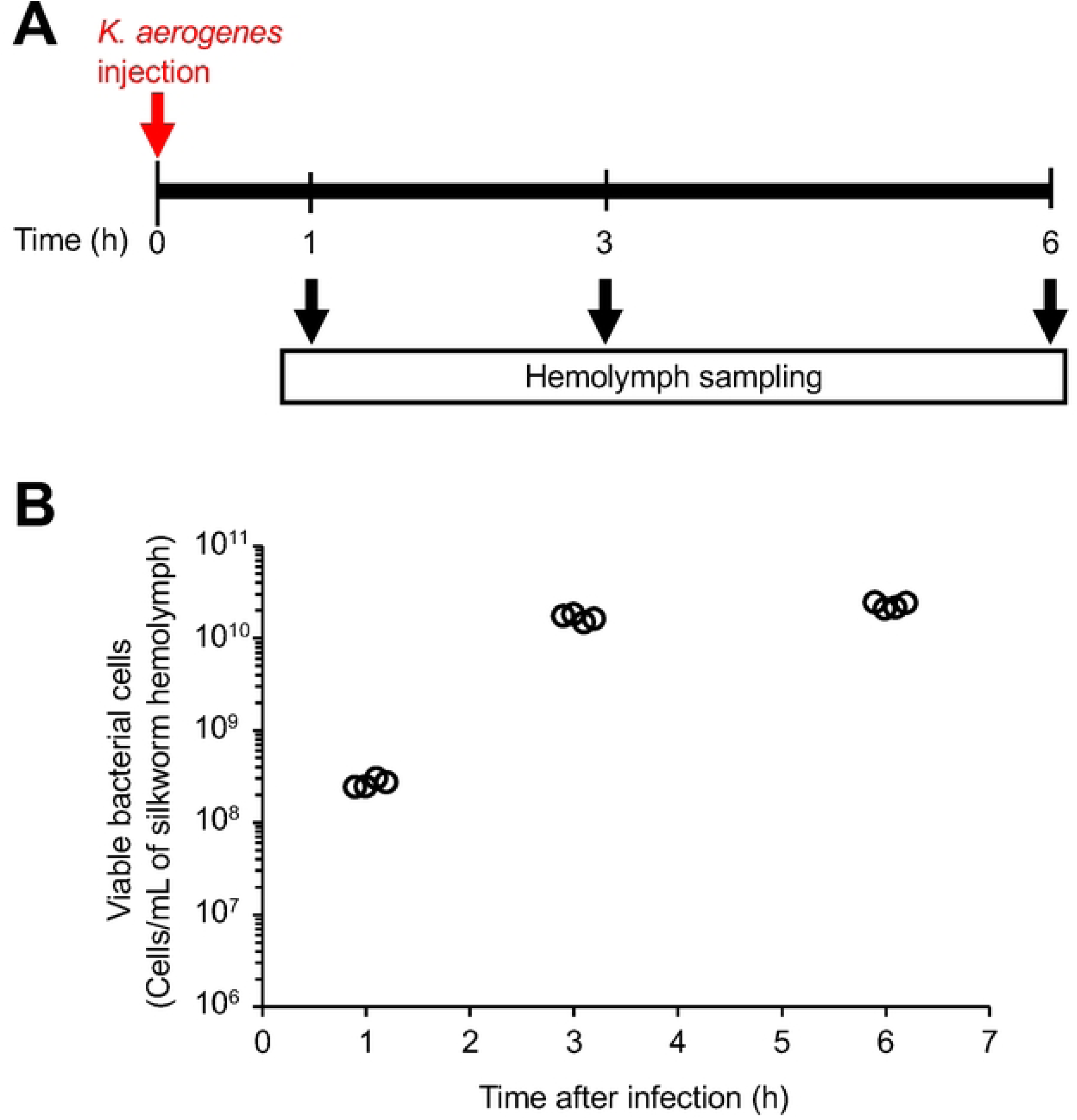
Increased viable cell number of *K. aerogenes* TUM25562 in the silkworm hemolymph. (**A**) Schematic representation of the infection experiment using silkworms. Silkworms were injected with a *K. aerogenes* TUM25562 suspension (7 x 10^6^ cells in 50 µl) and incubated at 37°C. Hemolymph was harvested from the silkworm at 1, 3, or 6 h post-infection. (**B**) Hemolymph was added to saline, and the solution was spread on an LB agar plate. The agar plate was incubated at 37°C for 1 day, and the colonies were counted. N = 4/group.

**Fig. 4.**
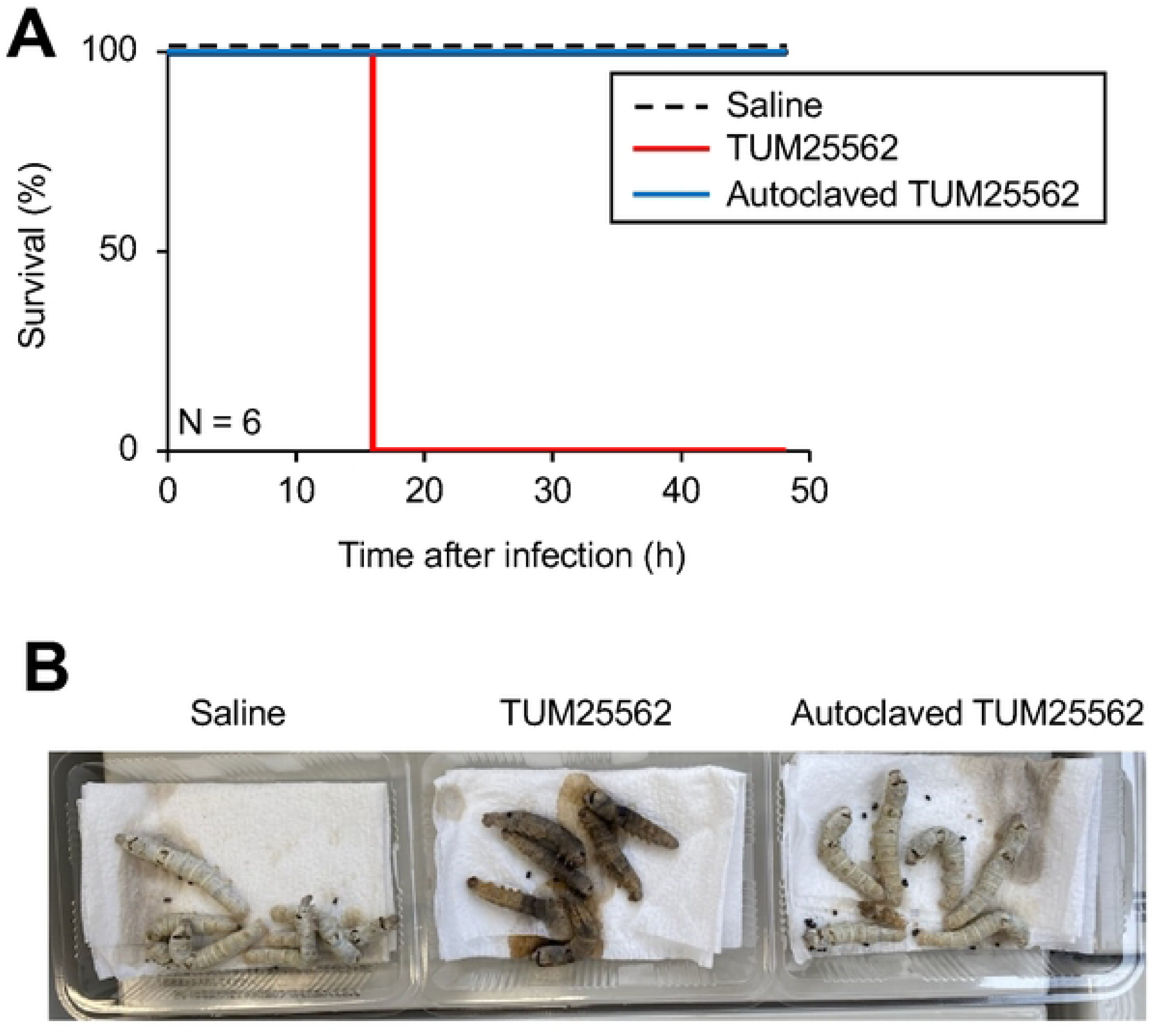
Effect of autoclave treatment of *K. aerogenes* TUM25562 on the killing ability against silkworms. (**A**) Time course of survival of silkworms injected with *K. aerogenes* TUM25562 or autoclave-treated *K. aerogenes* TUM25562. Saline (50 µL), *K. aerogenes* TUM25562 (A600 = 4) (50 µL), or autoclave-treated *K. aerogenes* TUM25562 (A600 = 4) (50 µL) was injected into the silkworms. After injection, the silkworms were incubated at 37°C for 2 days. Survival curves were generated using the Kaplan–Meier method. N = 6/group. (**B**) Photograph of silkworms at 24 h after injection.

**Table 2.** LD_50_ values of clinical *K. aerogenes* isolates in the silkworm infection model.

| *K. aerogenes* strains | LD<sub>50</sub> (x 10<sup>7</sup> cells/silkworm) |
| --- | --- |
| TUM25562 | 8.4 |
| TUM25564 | 18 |
| TUM25556 | 21 |

### Evaluation of clinically relevant antimicrobial doses in the silkworm infection model

Next, we investigated whether antimicrobial doses calculated from clinical weight-based guidelines were effective in the silkworm infection model. Based on the *Sanford Guide to Antimicrobial Therapy*, the recommended clinical dosage of MEPM is 40 mg/kg administered three times daily (120 mg/kg/day) [42,43]. Because the silkworms used in this study weighed approximately 2 g, the corresponding dose was calculated to be 80 µg/silkworm per administration (240 μg/day). The experimental design is shown in Fig. 5A. Administration of MEPM (80 µg/silkworm) three times daily did not prolong the survival of infected silkworms (Fig. 5B). Based on the *Sanford Guide to Antimicrobial Therapy*, the recommended clinical dosage of GM and AMK are 5–7.5 mg/kg once daily and 15 mg/kg once daily, respectively [43,44]. These doses correspond to 15 μg of GM and 30 μg of AMK per 2-g silkworm. Administration of GM (15 µg/silkworm) or AMK (30 µg/silkworm) did not prolong the survival of infected silkworms (Fig. 6). Based on the Johns Hopkins ABX Guide, the recommended clinical dosage of CFPM is 50 mg/kg administered three times daily (150 mg/kg/day) [43–45]. This dose corresponds to 100 μg/2-g silkworm. The experimental design is shown in Fig. 7A. Administration of CFPM (100 µg/silkworm) three times daily also failed to prolong the survival of infected silkworms (Fig. 7B). Together, these results indicate that antimicrobial doses calculated from clinical weight-based dosing guidelines did not improve survival in the silkworm infection model.

**Fig. 5.**
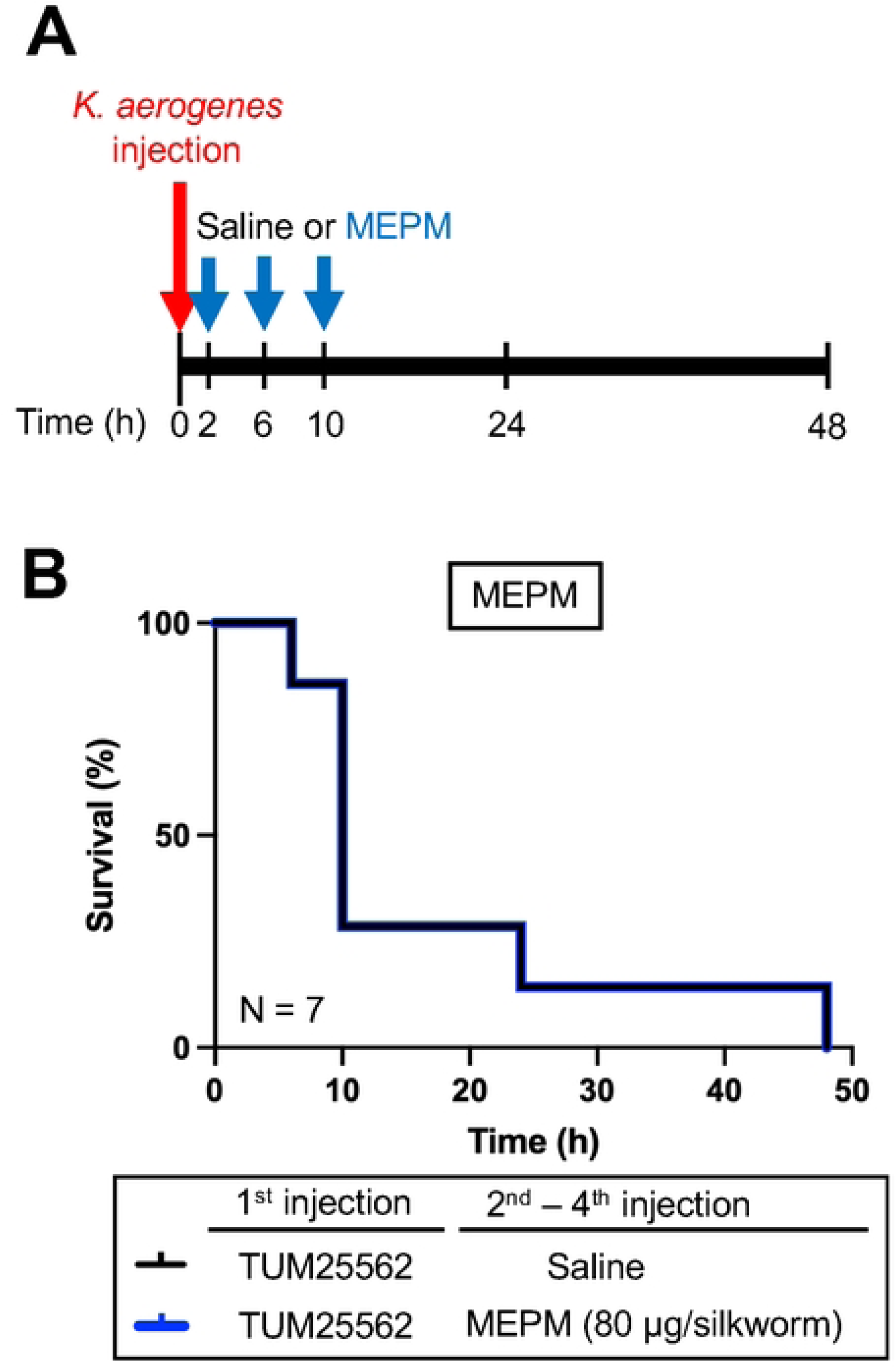
Effect of MPEM against silkworms infected with *K. aerogenes* TUM25562. (**A**) Experimental scheme of the MEPM administration against silkworms infected with *K. aerogenes* TUM25562. (**B**) Time course of survival of silkworms injected with *K. aerogenes* TUM25562. Saline (50 µL) or *K. aerogenes* TUM25562 (2 × 10⁸ cells/50 µL) was injected into the silkworms. The MEPM solution (80 µg/50 µL) or saline (50 µL) was administered three times a day at 2, 6, and 10 h after injection of *K. aerogenes* TUM25562. The silkworms were incubated at 37°C for 2 days. Survival curves were generated using the Kaplan–Meier method. N = 7/group.

**Fig. 6.**
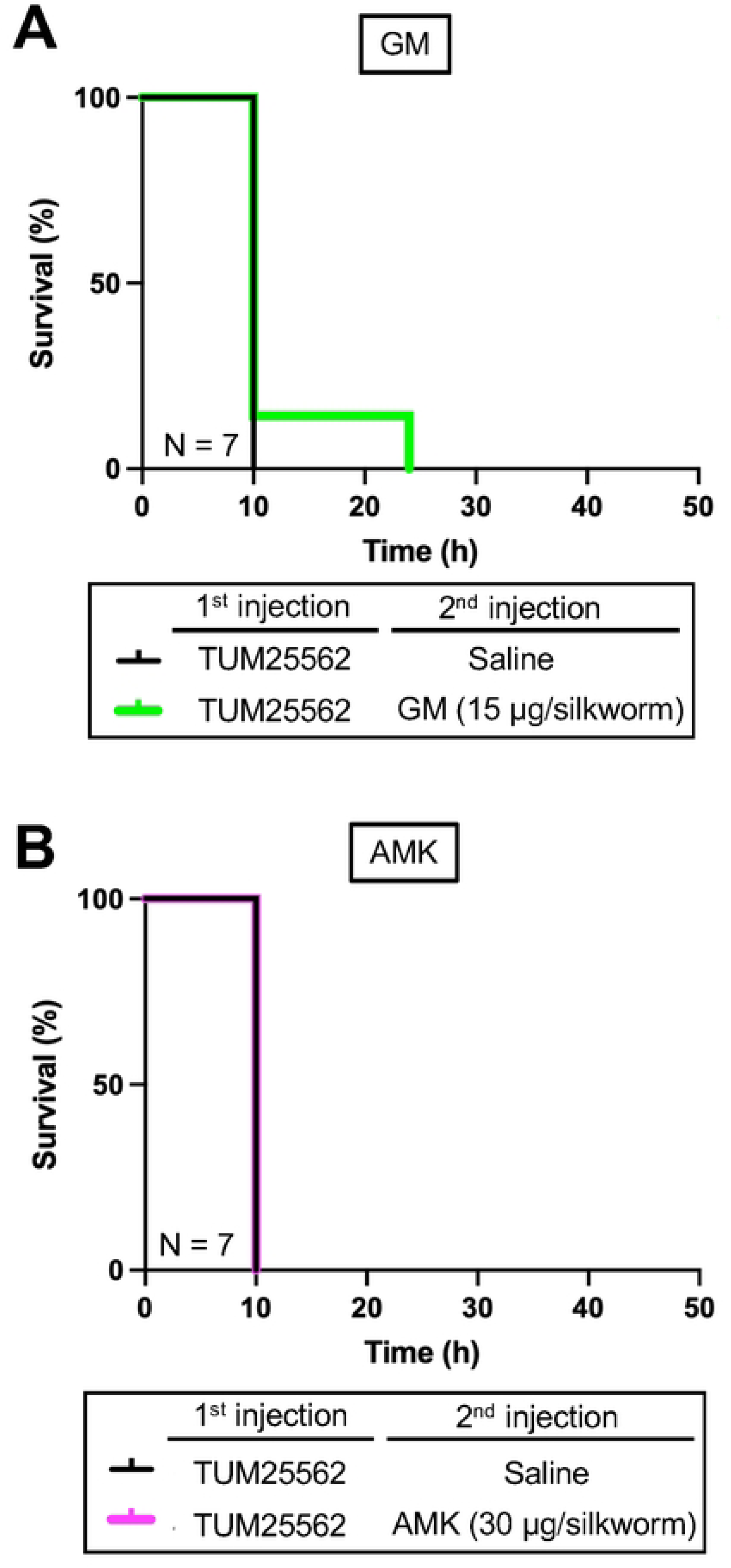
Effects of aminoglycosides against silkworms infected with *K. aerogenes* TUM25562. Saline (50 µL) or *K. aerogenes* TUM25562 (4 × 10⁸ cells/50 µL) was injected into the silkworms. The aminoglycoside solution [(**A**) GM: 15 µg/50 µL or (**B**) AMK: 30 µg/50 µL] or saline (50 µL) was administered 2 h after injection of *K. aerogenes* TUM25562. The silkworms were incubated at 37°C for 2 days. Survival curves were generated using the Kaplan–Meier method. N = 7/group.

**Fig. 7.**
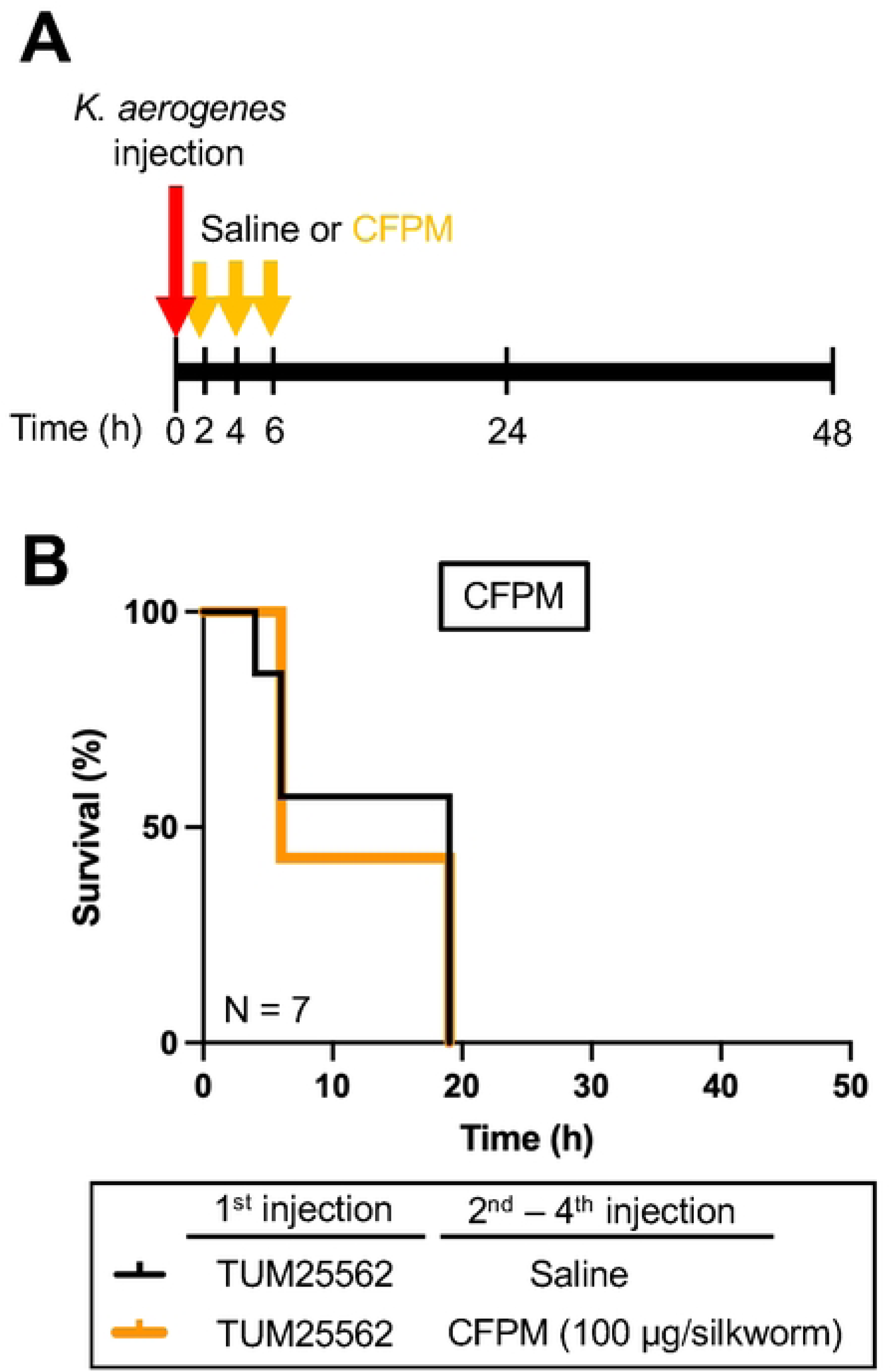
Effect of CFPM against silkworms infected with *K. aerogenes* TUM25562. (**A**) Experimental scheme of the CFPM administration against silkworms infected with *K. aerogenes* TUM25562. (**B**) Time course of survival of silkworms injected with *K. aerogenes* TUM25562. Saline (50 µL) or *K. aerogenes* TUM25562 (4 × 10⁸ cells/50 µL) was injected into the silkworms. The CFPM solution (100 µg/50 µL) or saline (50 µL) was administered three times a day at 2, 4, and 6 h after injection of *K. aerogenes* TUM25562. The silkworms were incubated at 37°C for 2 days. Survival curves were generated using the Kaplan–Meier method. N = 7/group.

### Therapeutic effects of high-dose antimicrobial therapy in the silkworm infection model

We next examined the therapeutic effects of high-dose antimicrobial administration in the silkworm infection model using *K. aerogenes* TUM25562. Administration of MEPM, GM, and AMK increased the number of surviving silkworms in a dose-dependent manner, whereas CFPM showed no dose-dependent therapeutic effect (Fig. 8). The ED_50_ values of MEPM, GM, AMK, and CFPM were 124, 199, 113, and >640 µg/silkworm, respectively (Table 3). Administration of MEPM and GM at four times the ED_50_ significantly prolonged the survival of silkworms infected with *K. aerogenes* TUM25562 (Fig. 9). We next evaluated the therapeutic effects of high-dose MEPM and GM in the silkworm infection model established using *K. aerogenes* TUM25564 and TUM25552. In silkworms infected with the *in vitro* MEPM-resistant isolate *K. aerogenes* TUM25564, administration of GM (800 µg/silkworm) prolonged survival, whereas MEPM (500 µg/silkworm) had no therapeutic effect (Fig. 10A). In contrast, in silkworms infected with the isolate TUM25556, which exhibited resistance to both MEPM and GM *in vitro*, administration of either MEPM (500 µg/silkworm) or GM (800 µg/silkworm) prolonged survival (Fig. 10B). These findings indicate that high-dose administration of MEPM or GM can improve the survival of silkworms infected with *K. aerogenes* TUM25562.

**Fig. 8.**
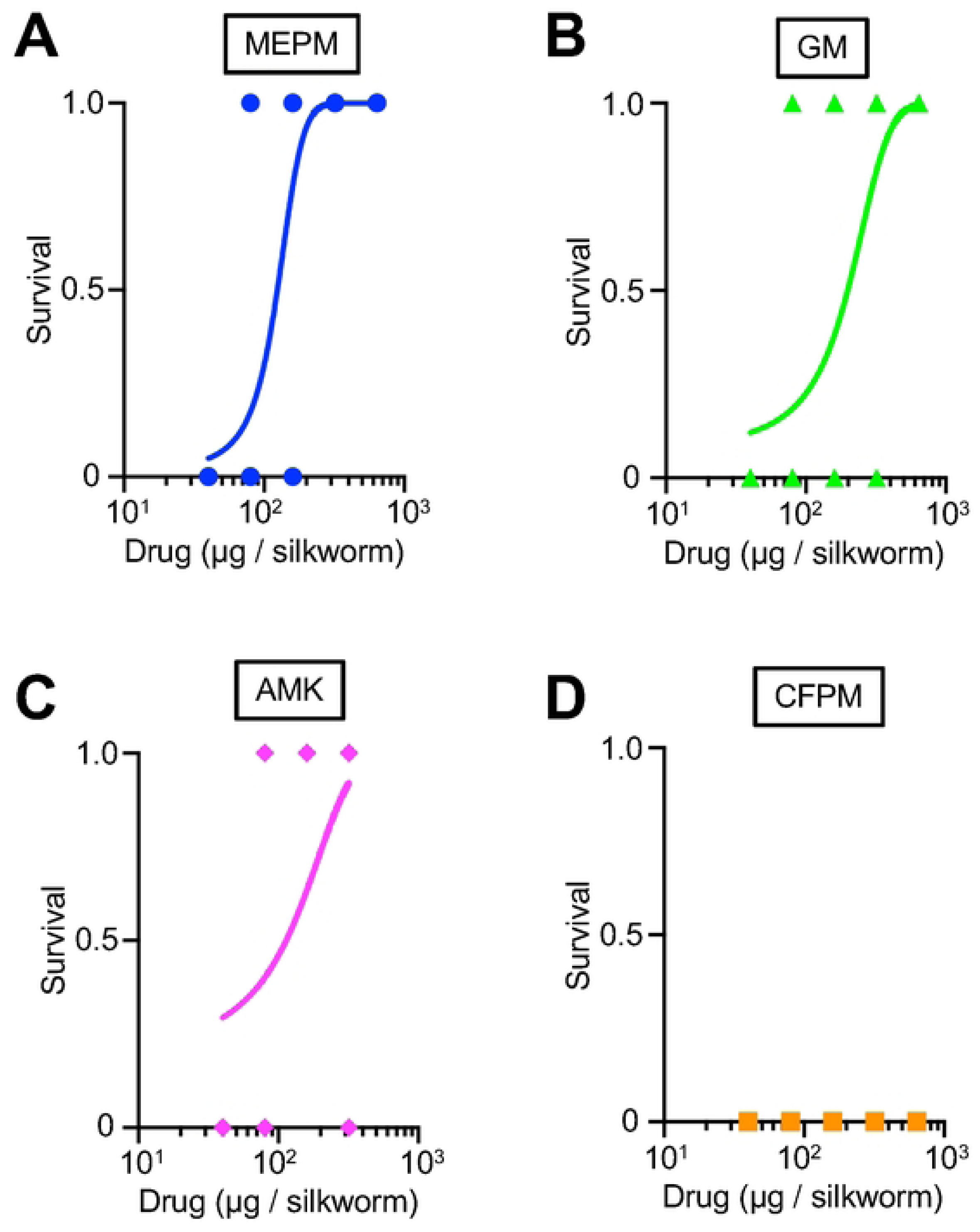
Dose-dependency of therapeutic effects of MEPM, GM, and AMK against silkworms infected with *K. aerogenes* TUM25562. *K. aerogenes* TUM25562 solution (4 × 10⁸ cells/50 µL) was injected into the silkworms. The antimicrobial agents (**A**) MEPM, (**B**) GM, (**C**) AMK, and (**D**) CFPM (40-320 µg/50 μL) were injected into the silkworm hemolymph at 2 h after infection. Silkworm survival at 24 h was monitored. Surviving and dead silkworms were scored as 1 and 0, respectively. A total of 20 silkworms were used per group. Curves were drawn using a simple logistic regression model.

**Fig. 9.**
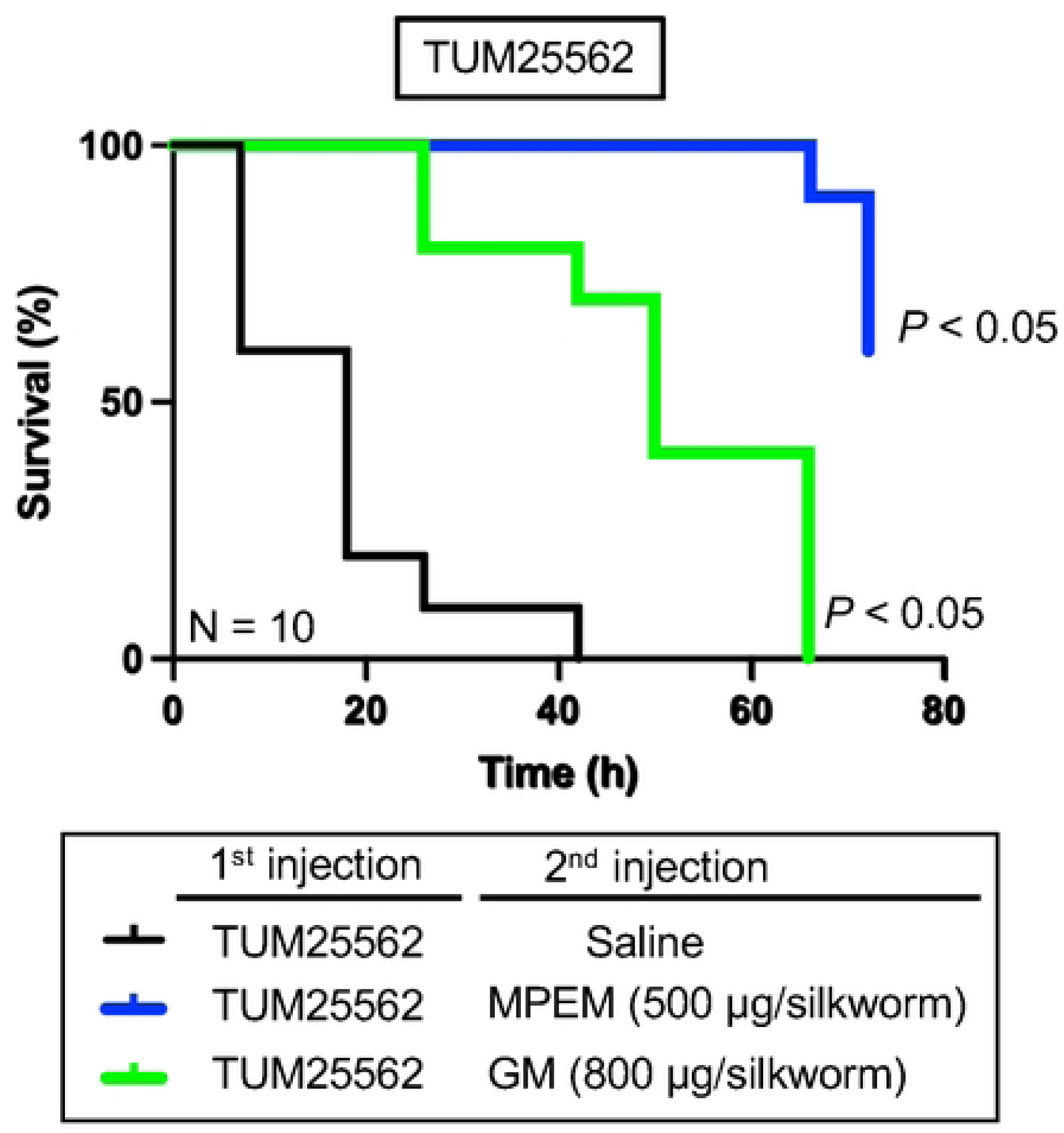
Therapeutic effects of high doses of MEPM and GM against silkworms infected with *K. aerogenes* TUM25562. *K. aerogenes* TUM25562 (4 × 10⁸ cells/50 µL) was injected into the silkworms. The antimicrobial solution [(**A**) MEPM: 500 µg/50 µL or (**B**) GM: 800 µg/50 µL] or saline (50 µL) was administered 2 h after injection of *K. aerogenes* TUM25562. The silkworms were incubated at 37°C for 3 days. Survival curves were generated using the Kaplan–Meier method. Statistical significance between the saline group and the drug-treated group was determined using the log-rank (Mantel–Cox) test. *P* < 0.05 was considered statistically significant. N = 10/group.

**Fig. 10.**
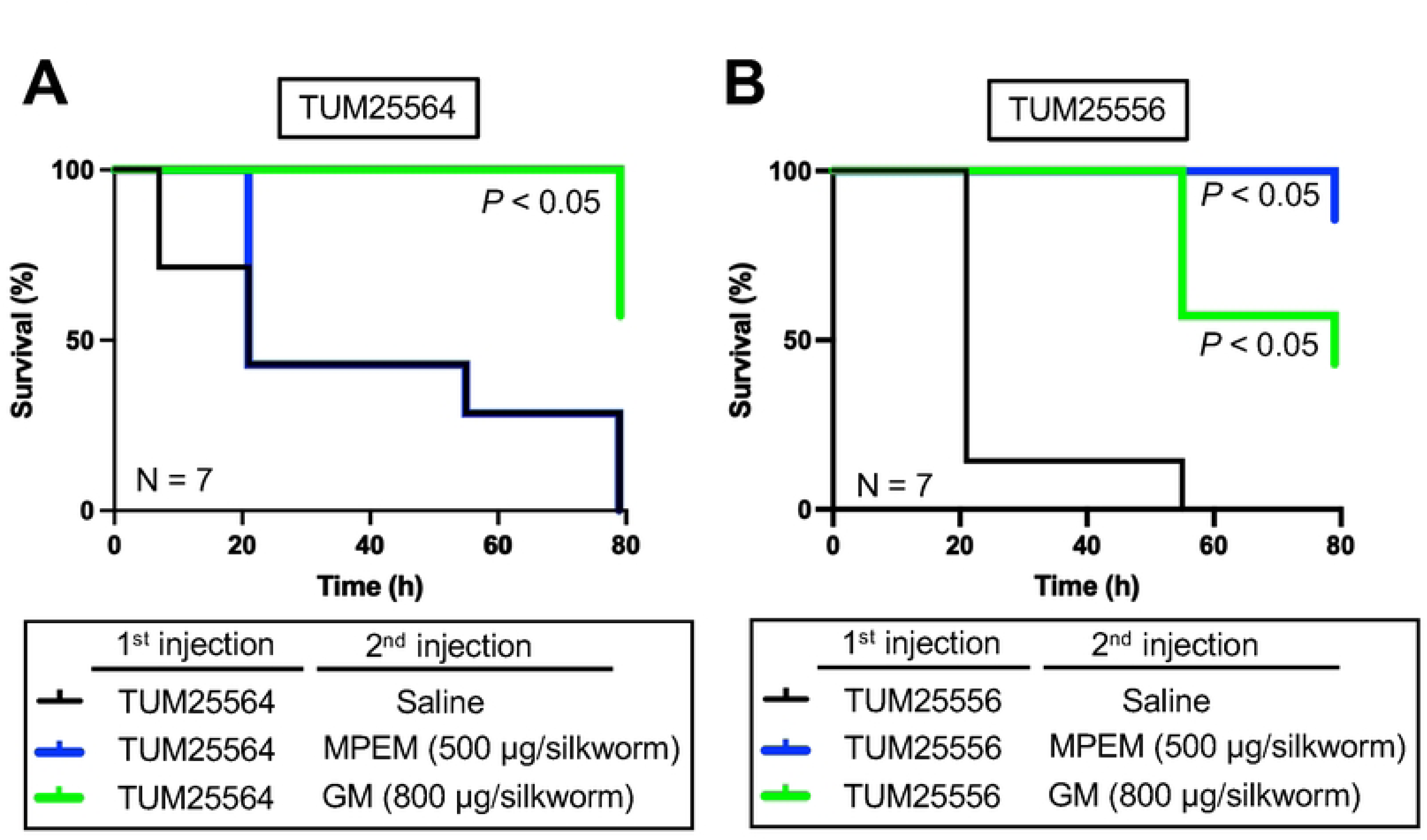
Therapeutic effects of high doses of MEPM and GM against silkworms infected with *K. aerogenes* TUM25564 or TUM25556. (**A**, **B**) *K. aerogenes* TUM25564 (4 × 10⁸ cells/50 µL) was injected into the silkworms. The antimicrobial solution [(**A**) MEPM: 500 µg/50 µL or (**B**) GM: 800 µg/50 µL] or saline (50 µL) was administered 2 h after injection of *K. aerogenes* TUM25564. The silkworms were incubated at 37°C for 3 days. Survival curves were generated using the Kaplan–Meier method. N = 7/group. (**C, D**) *K. aerogenes* TUM25556 (4 × 10⁸ cells/50 µL) was injected into the silkworms. The antimicrobial solution [(**C**) MEPM: 500 µg/50 µL or (**D**) GM: 800 µg/50 µL] or saline (50 µL) were administered 2 h after injection of *K. aerogenes* TUM25556. The silkworms were incubated at 37°C for 3 days. Survival curves were generated using the Kaplan–Meier method. Statistical significance between the saline group and the drug-treated group was determined using the log-rank (Mantel–Cox) test. *P* < 0.05 was considered statistically significant. N = 7/group.

**Table 3.**
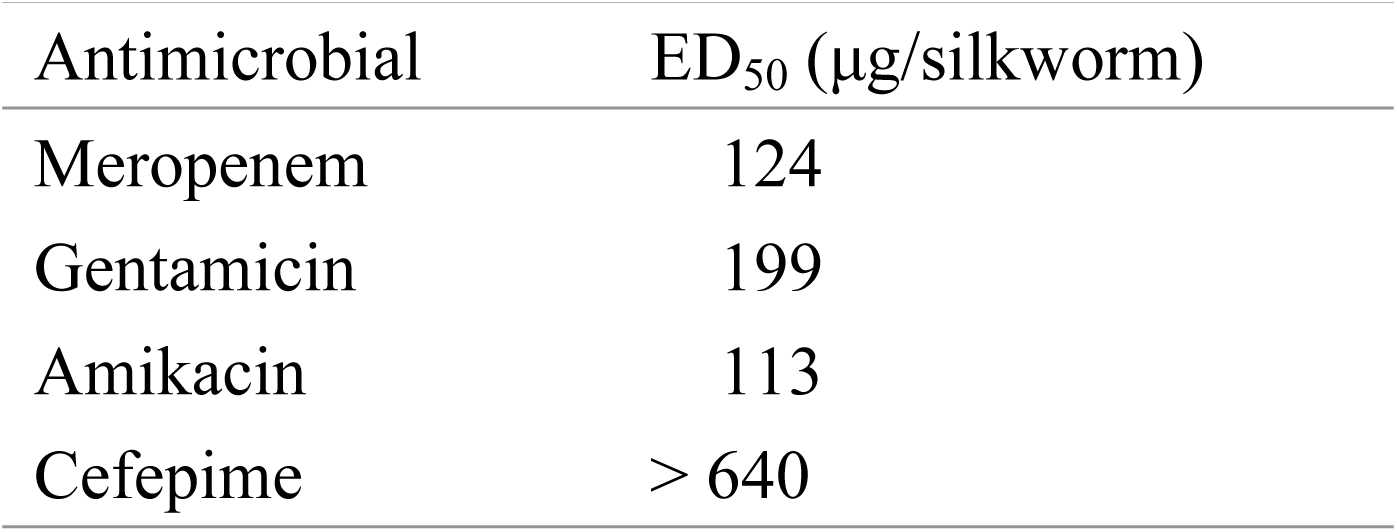
ED_50_ determination of antimicrobials against *K. aerogenes* TUM25562-infected silkworms.

| Antimicrobial | ED<sub>50</sub> (µg/silkworm) |
| --- | --- |
| Meropenem | 124 |
| Gentamicin | 199 |
| Amikacin | 113 |
| Cefepime | > 640 |

## Discussion

In this study, we isolated *K. aerogenes* strains from a patient and established a silkworm infection model using the clinical isolate TUM25562 to evaluate the therapeutic efficacy of antimicrobial agents. Guideline-recommended doses of MEPM, GM, AMK, and CFPM did not cure silkworms infected with *K. aerogenes* TUM25562. We therefore determined the ED_50_ values of these antimicrobial agents using the silkworm infection model. Administration of MEPM or GM at four times the ED_50_ significantly prolonged the survival of silkworms infected with *K. aerogenes* TUM25562. These findings suggest that a silkworm infection model using clinical *K. aerogenes* isolates may provide a practical *in vivo* platform for evaluating antimicrobial efficacy and determining effective antimicrobial doses, thereby supporting treatment selection for difficult-to-treat infections.

In the present clinical case, *K. aerogenes* isolates with progressively increased antimicrobial resistance were recovered during antimicrobial therapy. The initial isolate, TUM25562, exhibited reduced susceptibility to penicillin and cephalosporins but remained susceptible to MEPM and GM. During prolonged MEPM therapy, the MEPM-resistant isolate TUM25564 was recovered. Subsequently, after GM treatment, the GM-resistant isolate TUM25556 was recovered. These findings suggest that antimicrobial therapy may have contributed to the selection of drug-resistant *K. aerogenes* strains during treatment.

A silkworm infection model was established using *K. aerogenes* TUM25562 to evaluate bacterial virulence. Although a silkworm infection model for *K. aerogenes* has previously been reported [31], we established a model using the clinical isolate TUM25562 in this study. We first established experimental conditions under which more than half of the silkworms died within 24 h after injection of *K. aerogenes* TUM25562. Bacterial proliferation in the silkworm hemolymph was confirmed following infection, whereas silkworms injected with autoclave-treated *K. aerogenes* TUM25562 survived. These findings indicate that viable bacteria are required for silkworm killing and suggest that bacterial proliferation within the host contributes to lethality. Using this infection model, the LD_50_ values of *K. aerogenes* TUM25562, TUM25564, and TUM25556 were calculated 24 h after infection. Subsequent antimicrobial efficacy experiments were performed using an inoculum corresponding to four times the LD_50_.

Under dosing conditions based on the MEPM regimen used in the present clinical case, no therapeutic effect was observed in the silkworm infection model using *K. aerogenes* TUM25562. Similarly, when GM, AMK, and CFPM were administered at doses calculated from current clinical dosing guidelines, no therapeutic effect was observed. These findings suggest that antimicrobial doses derived from clinical dosing guidelines may not have been sufficient to achieve therapeutic efficacy in the silkworm infection model. On the other hand, *K. aerogenes* was susceptible to MEPM, GM, AMK, and CFPM *in vitro* based on MIC determinations. This discrepancy between the *in vivo* and *in vitro* results may reflect differences in antimicrobial pharmacokinetics or bacterial physiologic states within the host environment.

Therapeutic effects were observed in the silkworm infection model using *K. aerogenes* TUM25562 following the administration of high doses (four times the ED_50_) of MEPM or GM. Because the host environment can influence antimicrobial pharmacokinetics and bacterial susceptibility, we determined the ED_50_ values of antimicrobial agents in the silkworm infection model to identify therapeutically effective doses. The ED_50_ values of MEPM, GM, and AMK were successfully determined. On the other hand, no therapeutic effect was observed for CFPM; thus, its ED_50_ could not be determined. In addition, toxicity was observed at higher doses of AMK (Fig. S2). Therefore, subsequent analyses focused on the therapeutic effects of MEPM and GM. The ED_50_ values of MEPM and GM were approximately 1.5-fold and 13-fold higher, respectively, than the corresponding single doses recommended in the current clinical guidelines. Administration of MEPM or GM at four times the ED_50_ significantly prolonged the survival of silkworms infected with *K. aerogenes* TUM25562. These findings suggest that dose optimization using the silkworm infection model may help identify therapeutically effective antimicrobial doses for difficult-to-treat infections. In silkworms infected with *K. aerogenes* TUM25564, which exhibited resistance to MEPM *in vitro*, high-dose MEPM treatment was ineffective, whereas high-dose GM treatment prolonged survival. In silkworms infected with *K. aerogenes* TUM25556, which exhibited resistance to both MEPM and GM *in vitro*, treatment with either high-dose MEPM or GM prolonged survival. These findings indicate that antimicrobial activity observed in vivo does not necessarily correspond to *in vitro* susceptibility. Similar discrepancies were previously reported. For example, lysocin E, identified using a silkworm infection model, exhibits greater therapeutic activity *in vivo* in mice than predicted by its *in vitro* antimicrobial activity because its activity is enhanced by host-derived apolipoprotein A-I [46]. Likewise, ampicillin was therapeutically effective in a silkworm infection model using *K. aerogenes* strains that were resistant *in vitro* [31]. Clarifying the mechanisms underlying the discrepancies between *in vitro* and *in vivo* antimicrobial efficacy will be an important topic for future investigations.

## Conclusion

In this study, we established a silkworm infection model using clinical *K. aerogenes* isolates from a difficult-to-treat case and demonstrated its utility for experimentally evaluating the therapeutic efficacy of antimicrobial agents and determining effective antimicrobial doses. As invertebrate experimental animals, silkworms are inexpensive and raise few animal welfare concerns, making them well suited for large-scale infection experiments. In clinical cases where standard guideline-based antimicrobial therapy is ineffective, experimental simulations using silkworm infection models may provide additional information to support the selection of effective antimicrobial therapies.

## Acknowledgments

We thank Renta Endo, Momoka Matsumura, and Tomoya Sanbongi (Meiji Pharmaceutical University) for technical assistance in rearing the silkworms. We thank Dr. Kageto Yamada (Nihon University) and staff in the Microbiology Laboratory at Toho University Omori Medical Center for providing and analyzing the bacterial strains. We thank Dr. Kenichi Masumoto and Dr. Kohei Ogata (Department of Neonatology, Toho University Omori Medical Center) for obtaining clinical information. We also thank SciTechEdit International LLC (Highlands Ranch, CO, USA) for English language editing.

## Supporting information

**S1 File. The file includes Table S1, S2, Figs. S1, and S2.**

